# Pod indehiscence in common bean is associated to the fine regulation of *PvMYB26* and a non-functional abscission layer

**DOI:** 10.1101/2020.04.02.021972

**Authors:** Valerio Di Vittori, Elena Bitocchi, Monica Rodriguez, Saleh Alseekh, Elisa Bellucci, Laura Nanni, Tania Gioia, Stefania Marzario, Giuseppina Logozzo, Marzia Rossato, Concetta De Quattro, Maria L. Murgia, Juan José Ferreira, Ana Campa, Chunming Xu, Fabio Fiorani, Arun Sampathkumar, Anja Fröhlich, Giovanna Attene, Massimo Delledonne, Björn Usadel, Alisdair R. Fernie, Domenico Rau, Roberto Papa

## Abstract

In legumes, pod shattering occurs when mature pods dehisce along the sutures, and detachment of the valves promotes seed dispersal. In *Phaseolus vulgaris*, the major locus *qPD5.1-Pv* for pod indehiscence was identified recently. We developed a BC4/F4 introgression line population and narrowed the major locus down to a 22.5-kb region. Here, gene expression and a parallel histological analysis of dehiscent and indehiscent pods identified an *AtMYB26* orthologue as the best candidate for loss of pod shattering, on a genomic region ~11 kb downstream of the highest associated peak. Based on mapping and expression data, we propose early and fine up-regulation of *PvMYB26* in dehiscent pods. Detailed histological analysis establishes that pod indehiscence is associated to the lack of a functional abscission layer in the ventral sheath, and that the key anatomical modifications associated with pod shattering in common bean occur early during pod development. We finally propose that loss of pod shattering in legumes resulted from histological convergent evolution and that this is the result of selection at orthologous loci.

**One-sentence summary:** A non-functional abscission layer determines the loss of pod shattering; mapping data, and parallel gene expression and histological analysis support *PvMYB26* as the candidate gene for pod indehiscence.

## INTRODUCTION

Loss of seed shattering is a paradigmatic example of the changes that have occurred to crop plant traits compared to their wild progenitors, which collectively constitute the ‘domestication syndrome’ (Hammer 1984). In wild species, specialised seed-dispersal strategies are of fundamental importance for plant survival and fitness. Conversely, in domesticated forms, loss or reduction of seed shattering is desired to reduce yield losses.

Due to its complex evolutionary history, common bean (*Phaseolus vulgaris* L.) is an excellent model to study the domestication process (Bitocchi *et al.*, 2017), which included its parallel domestication in the Andes and Mesoamerica (Bitocchi *et al.*, 2013). In *P. vulgaris*, the dry beans are characterised by different degrees of pod shattering. These represent the majority of the domesticated pool (Gepts and Debouck 1991), where a limited level of pod shattering has been conserved to favour the threshing of the dry pods. Variations in the pod shattering intensity are also associated with the environmental conditions during maturation (e.g., humidity and temperature) (Parker *et al.*, 2020).

Secondary domestication events have resulted in the development of totally indehiscent snap-bean cultivars, with a dominance of the Andean gene pool among commercial snap beans (Wallace *et al.*, 2018). Snap beans are suitable for green pod production due to the low fibre content in the pod walls and sutures (i.e., the stringless type). Pioneering investigations into *Arabidopsis thaliana* have reconstructed the genetic pathways associated with its fruit differentiation and silique shattering, which provides a model of the mechanisms underlying seed dispersal for other crop species (for review, see Di Vittori *et al.*, 2019). In common bean, Koinange *et al.* (1996) identified the qualitative locus *St* on chromosome Pv02 for the presence of pod suture string. Their observation that pod fibre content correlates with pod shattering was confirmed by Murgia *et al.* (2017), who identified an association between the carbon and lignin contents and modulation of pod shattering. Nanni *et al.* (2011) and Gioia *et al.* (2013) identified the orthologous genes of *AtSHP* (Liljegren *et al.*, 2000) and *AtIND* (Liljegren *et al.*, 2004), respectively, in common bean, where *AtIND* was co-mapped with *St* (Koinange *et al.*, 1996). However, *PvSHP* and *PvIND* did not show any polymorphic sequences associated with occurrence of pod shattering (Nanni *et al.*, 2011, Gioia *et al.*, 2013). Recently, Rau *et al.* (2019) identified a major locus on chromosome Pv05 for pod indehiscence (*qPD5.1-Pv*), which was also confirmed by Parker *et al.* (2020). Rau *et al.* (2019) thus proposed a model in which at least three additional hypostatic loci on chromosomes Pv04, Pv05 and Pv09 are involved in modulation of pod shattering, with multifactorial inheritance of the trait previously suggested by Lamprecht (1932). The recent identification of a major locus for pod shattering in common bean (Rau *et al.*, 2019) and in cowpea (Lo *et al.*, 2018) in a syntenic region on chromosome Pv05 supports the occurrence of convergent molecular evolution in legume species. Moreover, Parker *et al.* (2020) suggested that the gene orthologous to *GmPDH1* in soybean (Funatsuki *et al.*, 2014) is also involved in the modulation of pod shattering in common bean.

In the present study, we developed a population of 1,197 BC4/F4 introgression lines (ILs) dedicated to pod shattering syndrome traits, with the aim to narrow down the major locus *qPD5.1-Pv*, and to promote recombination at QTLs for pod shattering. We also performed differential expression analysis at the transcriptome level (i.e., RNA-seq) between wild and domesticated pods, and at the major locus *qPD5.1-Pv* for target genes (i.e., qRT-PCR), using a comparison of indehiscent and highly dehiscent pods from near isogenic lines. The expression analysis for the putative structural genes of lignin biosynthesis and a parallel histological analysis of the indehiscent and dehiscent pods allow reconstruction of the main phenotypic events associated to the modulation of pod shattering, that occur early during pod development. Finally, through identification of orthologous genes, expression-analysis and selection signatures, we propose several candidate genes with potential roles in the modulation of pod shattering, both at the genome-wide level and at known QTLs.

## RESULTS

### Histological modifications underlying pod shattering in common bean

Lignification of the ventral and dorsal sheaths starts at 6 days after pod setting (DAP) for the pods of both the totally indehiscent variety Midas (Figure 1A, B; Supplemental Figure 1A, B) and the highly shattering IL 244A/1A (Figure 1C, D; Supplemental Figure 1C, D).

**Figure 1.**
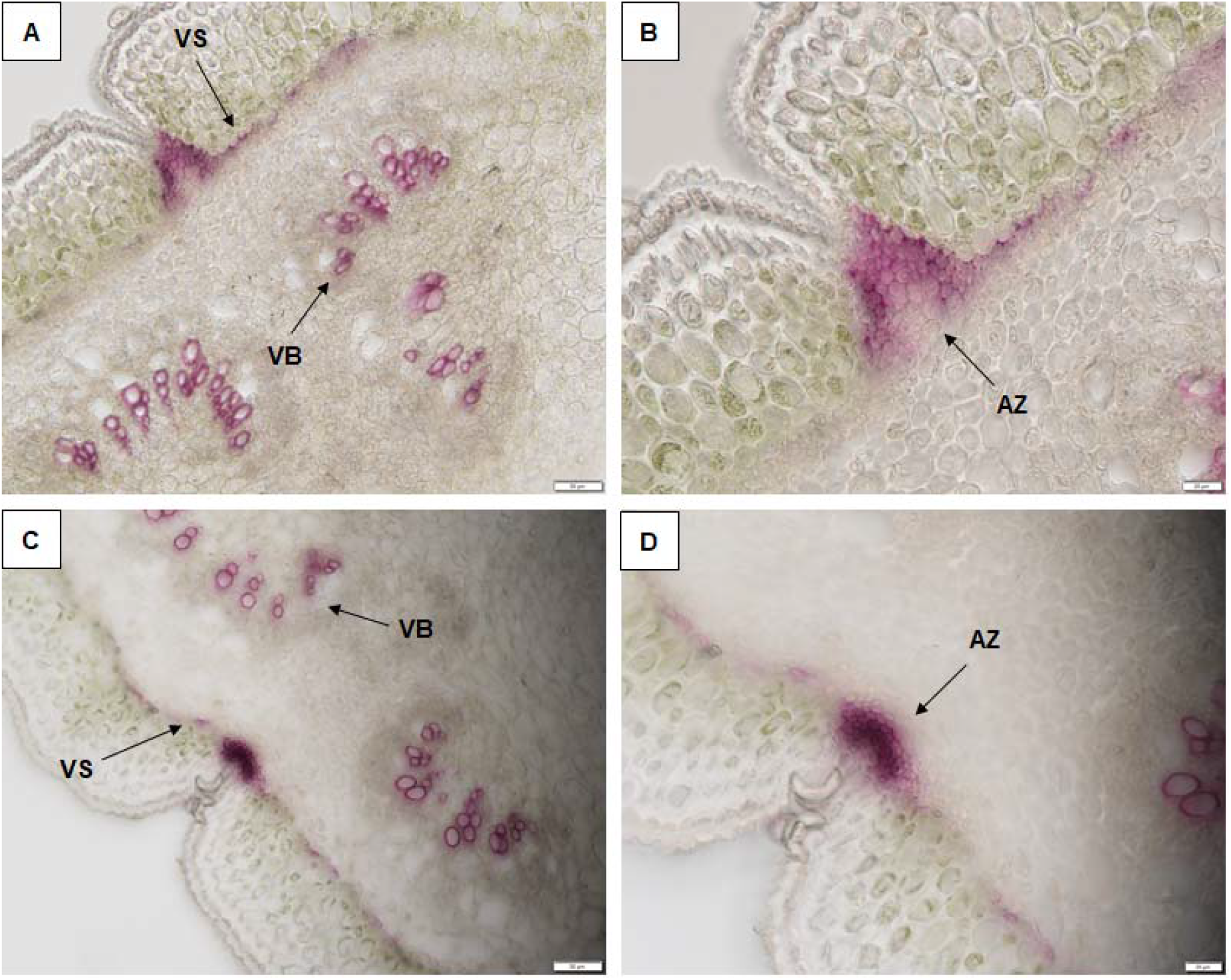
Analysis of lignification patterns in the ventral sheaths of 6-day-old pods of the totally indehiscent variety Midas (A, B) and the highly dehiscent IL 244A/1A (C, D). Cross-sections (section thickness, 30 μm) of pods after phloroglucinol staining for lignin. (B, D) Increased magnification from (A, C). Scale bars: 50 μm (A, C); 20 μm (B, D). VS, ventral sheath; VB, vascular bundles; AZ, abscission zone.

Higher lignification was seen here for both the ventral (Figure 1C, D) and the dorsal (Supplemental Figure 1C, D) sheaths of the highly shattering IL 244A/1A, compared to the corresponding tissues of the indehiscent genotype Midas (Figure 1A, B, Supplemental Figure 1A, B). Moreover, a different conformation of the ventral sheath was seen comparing these non-shattering and high-shattering pods. For 10-day-old pods (i.e., at 10 DAP), the lignification pattern of the ventral suture clearly differed between the totally indehiscent variety (Figure 2A, B) and the highly dehiscent recombinant inbred line (RIL) MG38 (Figure 2C, D) and IL 244A/1A (Figure 2E, F).

**Figure 2.**
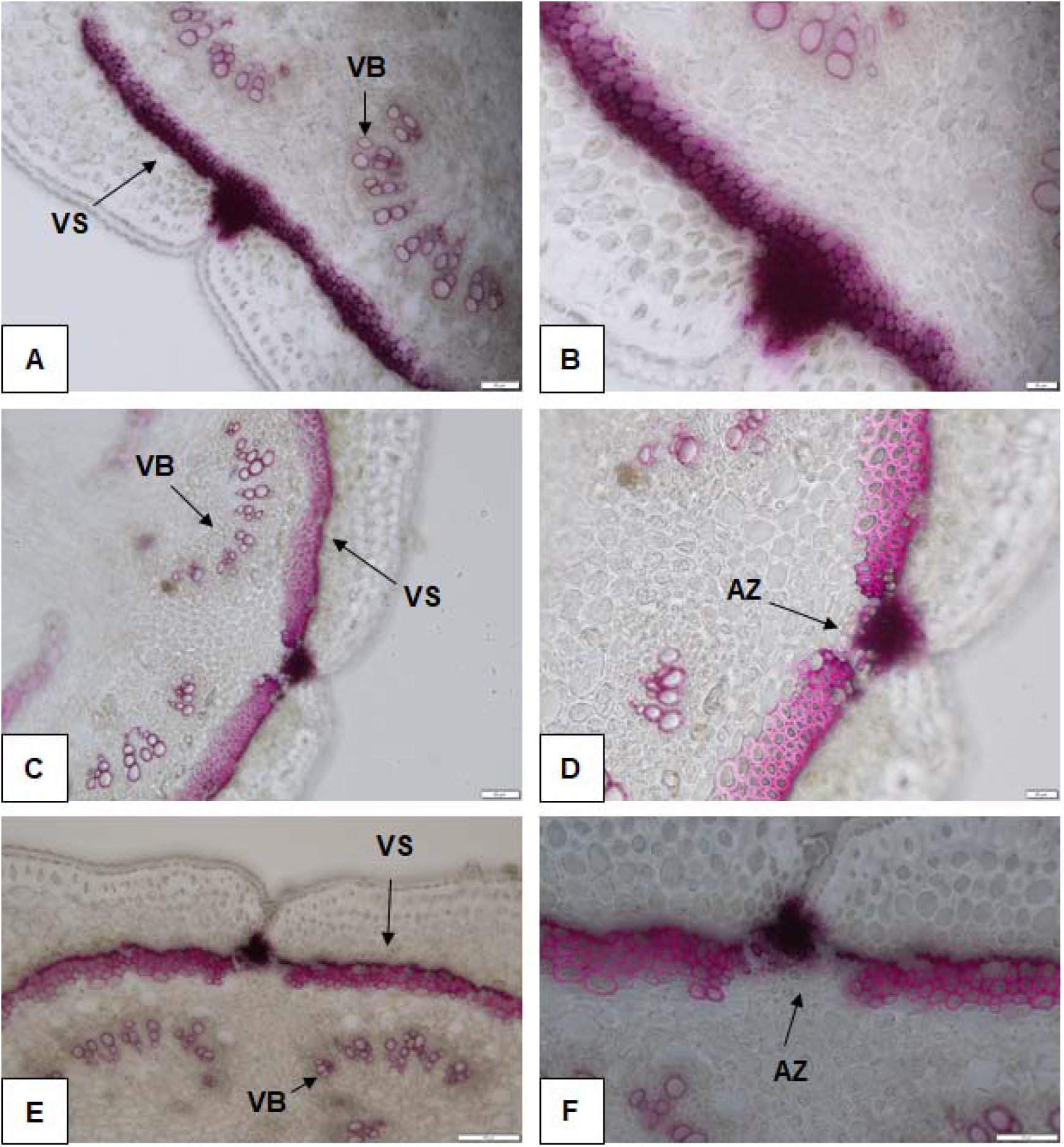
Analysis of lignification patterns in the ventral sheaths of 10-day-old pods of the totally indehiscent variety Midas (A, B) and the highly dehiscent RIL MG38 (C, D) and IL 244A/1A (E, F). Cross-sections (section thickness, 30 μm) of pods after phloroglucinol staining for lignin. (B, D, F) Increased magnification from (A, C, E). Scale bars: 50 μm (A, C, F); 20 μm (B, D); 100 μm (E). VS, ventral sheath; VB, vascular bundles; AZ, abscission zone.

A few layers of cells were lignified in the abscission zone of the non-shattering type (Figure 2B), compared to the equivalent tissue of the highly shattering lines (Figure 2D, F), which lacked lignification. This modification is potentially involved in prevention of pod opening. The walls of the cells that surrounded the abscission zone in the ventral sheath were heavily thickened in the highly shattering pods (Figure 2D, F), compared to the equivalent cells of the totally indehiscent pods (Figure 2B). This might increase the mechanical tension within the ventral suture, to thus promote pod shattering. Moreover, at 10 DAP, the highly shattering pods showed an internal lignified valve layer (Supplemental Figure 2B, C), which was not seen for the indehiscent pods of the variety Midas (Supplemental Figure 2A). At 14 DAP, the degree of lignification of the ventral suture, and both the ventral sheath and the abscission zone conformations strongly differed between the indehiscent variety Midas (Supplemental Figure 3A, B) and the highly shattering RIL MG38 (Supplemental Figure 3C, D) and IL 244A/1A (Supplemental Figure 3E, F). The histological conformation of mature pods at 30 DAP is presented in Figure 3.

**Figure 3.**
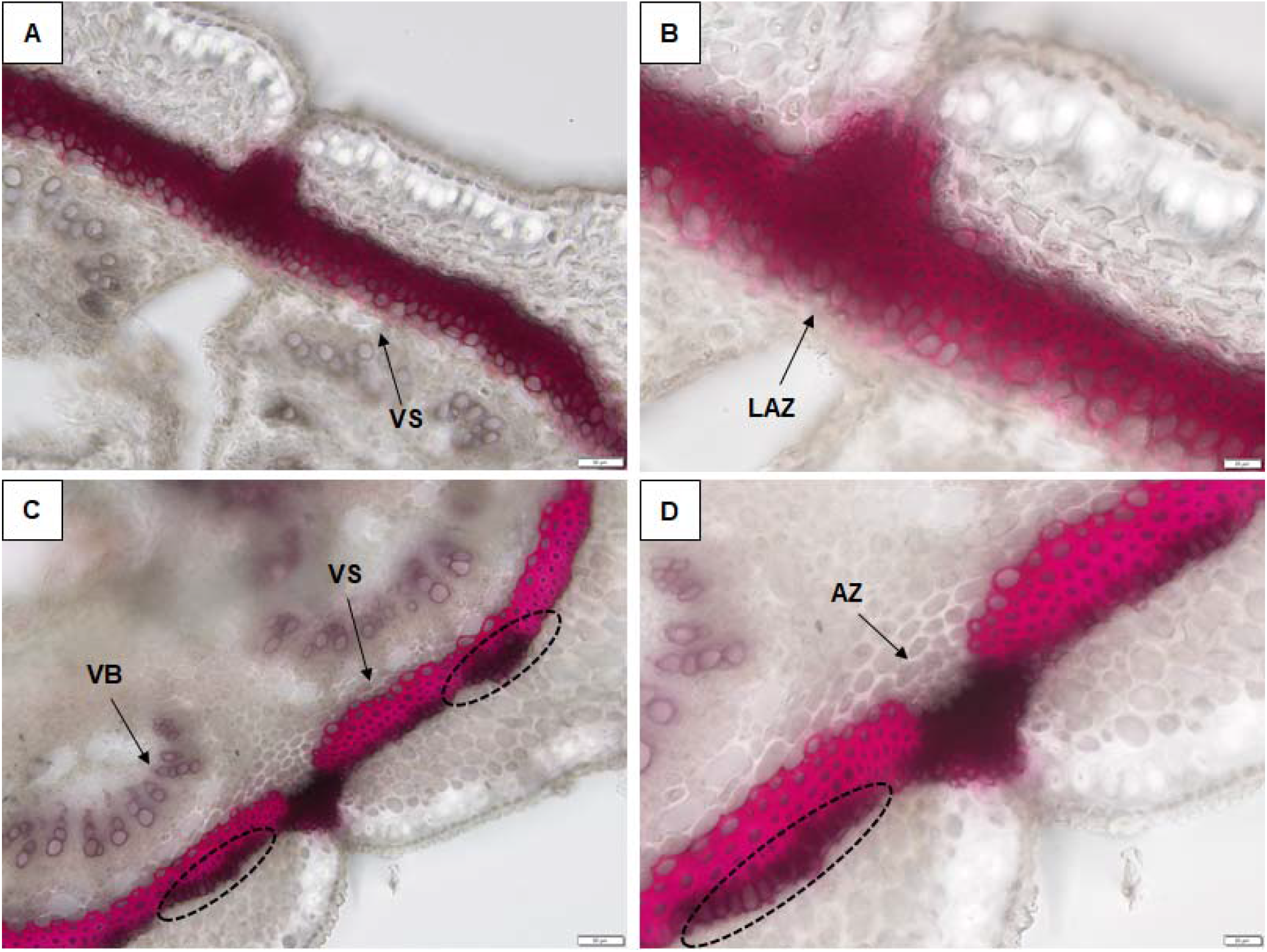
Analysis of lignification patterns of the ventral sheaths in 30-day-old pods (i.e., mature pods) of the totally indehiscent variety Midas (A, B) and the highly dehiscent IL 038A/2A2 (C, D). Cross-sections (section thickness, 50 μm) of the ventral suture after phloroglucinol staining for lignin. (B, D) Increased magnification from (A, C). Scale bars: 50 μm (A, C); 20 μm (B, D). VS, ventral sheath; VB, vascular bundles; AZ, abscission zone; LAZ, lignified abscission zone. (C, D) Dotted ellipses, lignification areas with no strong cell wall thickening along the ventral sheath.

In the region where the pods open at maturity (i.e., the abscission zone), in the highly shattering type, there were five layers of cells that completely lacked lignification of the cell walls (Figure 3D), compared to the lignification of the equivalent cells for the totally indehiscent pods (Figure 3B). We therefore suggest that the non-functional abscission layer is responsible for the loss of pod shattering in common bean. The cell walls were heavily thickened in the ventral sheath of the high-shattering pods (Figure 3D), compared to those of the ventral sheath of the indehiscent pods (Figure 3B). The lumen of the cells also appeared to be almost occluded in some of the cells of the high-shattering pod sheaths. Interestingly, there were a few layers of lignified, but not heavily thickened, cells across the ventral sheath of the mature dehiscent pods (Figure 3C, D, dashed ellipses). It is possible that different degrees of wall thickening along the sutures is required to create the mechanical tension needed for pod shattering and/or pod twisting.

### Segregation of pod shattering

Phenotyping for pod shattering on 100 lines from six BC4/F1 families revealed uniformity in F1 for the presence of pod shattering. Phenotyping of a subset of 509 BC4/F2 lines, from the first planting and that uniformly reached the maturation, identified 386 and 120 dehiscent and indehiscent plants, which fits the 3:1 expected ratio of a Mendelian trait (χ^2^ = 0.45) (Supplemental Table 1). The expected segregation ratio of a Mendelian trait was also observed when each of the BC4/F2 subpopulations were analysed separately (Supplemental Table 2), and for a subset of lines from the BC4/F3 population that produced enough pods for a reliable post-harvest phenotyping of pod shattering (356 putative dehiscent *vs* 193 putative indehiscent lines) (χ^2^ = 1.28) (Supplemental Table 3). Moreover, 354 BC4/F2 dehiscent ILs showed pod twisting to different degrees (classed as: 1% to <10%; ≥10% to <24%; ≥24%; Supplemental Table 1), while 32 dehiscent lines did not show any twisting; assuming the action of duplicated and independent genes with cumulative effects, this fits to a 15:1 twisting/ non-twisting ratio (χ^2^ = 2.74).

Due to the high correlation that was observed here between the field and post-harvest phenotyping of the BC4/F2 population (r = 0.81; p = 7.33 * 10^−118^), the post-harvest evaluation was also integrated into the subsequent analysis. In all, 1,197 BC4/F4 ILs were phenotyped for pod shattering in the field and/or after harvesting. When the field and post-harvest phenotypes were combined (i.e., defined as the ‘SH y/n’ [pod shattering, yes/no] trait), 940, 11 and 243 ILs were classified as dehiscent, intermediate and indehiscent, respectively (Supplemental Table 4). Overall, 721 F3 families were represented at the beginning of the BC4/F4 field experiment (i.e., 2,230 BC/F4 seeds were sown from 721 F3 plants), from which 502 F3 families produced BC4/F4 progenies. Of these, and as expected, 95 indehiscent F3 lines gave complete indehiscent F4 progeny, while segregation was still observed within 55 F3 families.

### Genome-wide association study for pod shattering and fine mapping of the major locus *qPD5.1-Pv*

A genome-wide association study (GWAS) for pod shattering was performed using a dataset of 19,420 single-nucleotide polymorphisms (SNPs) from genotype-by-sequencing (GBS) analysis, which were identified across 1,196 BC4/F4 ILs (Supplemental Figure 4). GWAS for the trait defined as ‘SH y/n’ (dehiscent *vs* indehiscent lines) identified a major locus for occurrence of pod shattering at the end of chromosome 5 (*qPD5.1-Pv*) (Figure 4); here, 52 SNPs showed association (-log_10_p >6) with the presence/ absence of pod shattering in the interval between the S5_38322754 and S5_39384267 markers.

**Figure 4.**
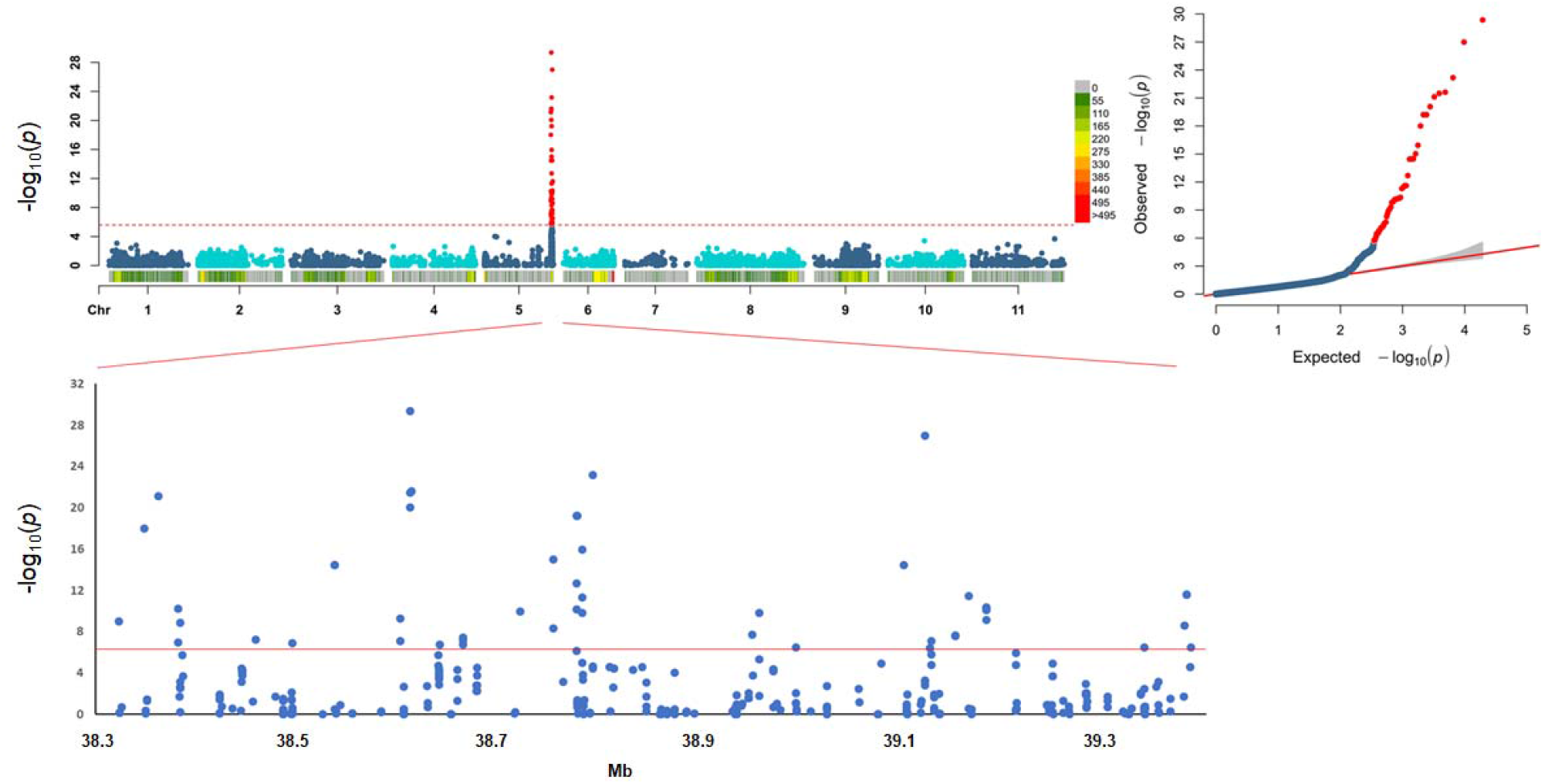
Genome-wide association study for occurrence of pod shattering. Top left: Manhattan plot to show the associations between 52 SNP markers (red dots on distal region of chromosome Pv05) and the SH y/n trait (dehiscent *vs* indehiscent lines). Dashed red line, fixed threshold of significance for the 19,420 SNP markers physically distributed across the 11 common bean chromosomes. Top right: QQplot of the distribution of the observed p values compared to the expected distribution. Bottom: Expanded major QTL on the distal part of chromosome Pv05, defining the significance of the SNP markers from 38.3 to 39.4 Mb on chromosome Pv05.

The major locus *qPD5.1-Pv* was also in the association for the following mapping analyses: when 18 ILs for which the phenotype score was not clearly assigned were removed (see Supplemental Table 4) (Supplemental Figure 5A); when the ‘SH y/n’ trait that included plants with an intermediate phenotype was used (Supplemental Figure 5B); when the presence/ absence of pod shattering was only from the field phenotyping (Supplemental Figure 5C); when the post-harvest phenotype was used (putative dehiscent *vs* putative indehiscent lines; Supplemental Figure 5D); when all of the phenotypic classes from the post-harvest evaluation were used (quantitative score; Supplemental Figure 5E); and when the percentage of twisting pods per plant was used (field evaluation; Supplemental Figure 5F). These GWAS data are summarised in the Supplemental Table 5, while Supplemental Figure 6 shows the expanded major QTL for all of these mapping strategies. A few recurrent highly associated SNPs were identified within the major locus (Figure 4; Supplemental Figure 6; and Supplemental Table 5). These identified three genomic regions around 38.61 Mb, 38.79 Mb and 39.12 Mb on chromosome Pv05. In particular, S5_38611412 was among the best associated SNPs for all of the mappings, with a few surrounding SNPs with high p values (Supplemental Table 5; Supplemental Figure 6). After narrowing the QTL to a 22.5 kb surrounding region (from S5_38605293 to S5_38627793), a few candidates were identified, among which there was a protein kinase (Phvul.005G157300), a phospholipid-transporting ATPase (Phvul.005G157400; with the highest associated SNP S5_38611412) and a nucleotidase (Phvul.005G157500). The main peak was located ~11 kb before a MYB26 transcription factor (Phvul.005G157600), the orthologue that is involved in anther dehiscence and secondary cell-wall differentiation in *A. thaliana* (Yang *et al.*, 2007). Moreover, a cluster of lipoxygenase genes were located on a tightly associated genomic region, from ~48 kb to ~17 kb upstream of the main peak (Phvul.005G156700, Phvul.005G156800, Phvul.005G156900, Phvul.005G157000).

### Identification of candidate genes for pod shattering and gene-expression analysis

The candidate genes were identified based on the annotation, the function of orthologues in legume species and *A. thaliana*, the differential expression analysis using RNA-seq data between wild and domesticated pods, the differential expression at the target candidate genes for the major locus *qPD5.1-Pv* (qRT-PCR) in a comparison of near isogenic lines, and the evidence of selection signatures from Schmutz *et al.* (2014) and Bellucci *et al.* (2014).

#### *Candidate genes at the major locus* qPD5.1-Pv

Supplemental Data Set 1 summarises the genes within the major locus *qPD5.1-Pv*, along with their differential expression and selection signatures. Overall, *qPD5.1-Pv* contains 128 genes, of which 29 were differentially expressed (from RNA-seq data), and 15 were under selection in the Mesoamerican gene pool, according to Schmutz *et al.* (2014) and/or Bellucci *et al.* (2014). Four genes were both differentially expressed and under selection.

Located ~11 kb downstream to the most significant peak, Phvul.005G157600 is orthologous to *AtMYB26* (Yang *et al.*, 2007). Phvul.005G157600 expression was up-regulated in 5-day-old dehiscent pods (i.e., Midas *vs* G12873), and down-regulated in G12873 dehiscent pods at 10 DAP (Supplemental Data Set 1; row 50). Down-regulation of *PvMYB26* expression was also seen for the comparison of Mesoamerican domesticated and wild (MD *vs* MW) at 5 DAP. Moreover, two genes located downstream to *PvMYB26* (Phvul.005G157700, Phvul.005G157800) on the physical map showed signatures of selection.

Within the highest associated region to which *qPD5.1-Pv* was narrowed down (S5_38605293:S5_38627793), Phvul.005G157400 and Phvul.005G157500 did not show differential expression or selection signatures, while no reads were mapped (i.e., RNA-seq) on Phvul.005G157300 in any of the samples (Supplemental Data Set 1; rows 47-49). In addition, *qPD5.1-Pv* contained a cluster of three differentially expressed linoleate 9S-lipoxygenase genes (Phvul.005G156700, Phvul.005G156900, Phvul.005G157000; Supplemental Data Set 1; rows 41, 43, 44) that were located upstream (from ~48 to ~17 kb) of the highest associated peak for pod indehiscence. In more detail: Phvul.005G156700 was down-regulated for Midas *versus* G12873 and Andean domesticated snap bean (AD_Snap) *versus* Andean wild (AW) at 10 DAP; Phvul.005G156900 expression was up-regulated for Midas *versus* G12873 at 10 DAP; while Phvul.005G157000 was down-regulated for the totally indehiscent pods (Midas *vs* G12873) at 10 DAP, and also showed signatures of selection in the Mesoamerican gene pool. In the region that surrounds SNP S5_39120955, which was also highly associated to occurrence of pod shattering (see Supplemental Table 5), there was a cluster of leucine-rich repeat (LRR) coding genes. In particular, Phvul.005G163800 and Phvul.005G163901 (Supplemental Data Set 1; rows 108, 110), which are both annotated as LRR-protein-kinase related, show differential expression for AD_Snap *versus* AD, AD *versus* AW, and MD *versus* MW at 5 DAP (Phvul.005G163800), and for Midas *versus* G12873 at 10 DAP (Phvul.005G163901). SNP S5_38792327 was also one of the best associated SNPs at the major locus *qPD5.1-Pv* (see Supplemental Table 5), and it was located within a fatty acid omega-hydroxy dehydrogenase (Phvul.005G159400; Supplemental Data Set 1; row 69), which, however, did not show selection signatures or significant differential expression. Finally, Phvul.005G164800 showed higher expression in indehiscent pods of Midas at 5 DAP and 10 DAP, compared to G12873 (Supplemental Data Set 1; row 119), and it was annotated as ZINC FINGER FYVE-DOMAIN-CONTAINING PROTEIN.

#### Candidate genes with a putative function in pod shattering based on their orthologues

Orthologous genes in common bean that in other species have pivotal roles in modulation of pod shattering, cell-wall modifications and putative pod-shattering-related functions were identified. These orthologous genes are reported in Supplemental Data Set 2, along with the results of the differential expression analysis (i.e., RNA-seq) and the signatures of selection.

First, we consider as promising candidates the orthologous genes located close to the known QTLs for pod shattering. On chromosome Pv02, Phvul.002G271000 (*PvIND*; [Gioia *et al.*, 2013]) is orthologous to *AtIND* (Liljegren *et al.*, 2004), and it was highly expressed in the snap-bean group compared to AW at 10 DAP (Supplemental Data Set 2; row 28); moreover, close to *PvIND*, we identified the NAC transcription factor Phvul.002G271700 (orthologous to NAC082). Both Phvul.002G271000 and Phvul.002G271700 map to the *St* locus (Koinange *et al.*, 1996). On chromosome Pv03, Phvul.003G252100 is orthologous to *Glycine max PDH1* (Funatsuki *et al.*, 2014), which was recently proposed as a candidate for modulation of pod shattering in common bean (Parker *et al.*, 2020); here, Phvul.003G252100 was up-regulated for Midas *versus* G12873 at 5 DAP and 10 DAP, and down-regulated for AD *versus* AW at 5 DAP, and MD *versus* MW at 10 DAP, with a signature of selection in the Andean gene pool (Supplemental Data Set 2; row 45). On chromosome Pv04, Phvul.004G144900 is orthologous to the MYB52 transcription factor, which maps to a region associated with modulation of pod shattering (Rau *et al.*, 2019); here, Phvul.004G144900 was less expressed for AD_Snap *versus* AW and MD *versus* MW, both at 10 DAP (Supplemental Data Set 2; row 50). Moreover, ~660 kb downstream, Phvul.004G150600 is a PIN family member, and thus putatively involved in correct regulation of auxin efflux. Phvul.004G150600 showed higher expression for indehiscent pods (Midas *vs* G12873) at 5 DAP, with a signature of selection (Supplemental Data Set; row 51). On chromosome Pv09, close to the significant SNP for shattering modulation at ~30 Mb that was identified by Rau *et al.* (2019), and within the QTL identified also by Parker *et al*. (2020), Phvul.009G203400 is the orthologous to *AtFUL* (Gu et *al.*, 1998); interestingly, Phvul.009G203400 shows parallel selection between the gene pools (Schmutz *et al.*, 2014), and congruently across different studies (Schmutz *et al.*, 2014 and Bellucci *et al.*, 2014) (Supplemental Data Set 2; row 93). In the same region, two physically close genes, Phvul.009G205100 and Phvul.009G205200, are orthologous to *Cesa7*, and they showed selection signatures. Moreover, Phvul.009G205100 was less expressed in the domesticated pods (Supplemental Data Set 2; rows 94, 95).

Here, we also identified potential candidates at the genome-wide level based on their orthology with genes with well-described functions in the modulation of seed dispersal and/or fruit development in other species, and because they showed signatures of selection and/or interesting differential expression patterns. Those that can be highlighted are: Phvul.002G294800, as orthologous to *GmPDH1*; Phvul.003G166100 and Phvul.011G100300, as putative orthologous to *Sh1*; Phvul.003G182700 and Phvul.003G281000, as orthologous to *AtFUL*; Phvul.007G100500, as putative orthologous to *Shattering4*; Phvul.008G114300 and Phvul.010G011900, as orthologous to *Replumless*, *SH5* and *qSH1*; and in particular, Phvul.010G118700, as orthologous to *NST1* and *GmSHAT1-5* (Supplemental Data Set 2). These data suggest that these genes might share a conserved pod-shattering-related function. Moreover, an *AtMYB26* orthologue on chromosome Pv10, Phvul.010G137500, was under-expressed in the AD and AD_Snap pods, compared to the wild pods at 5 DAP, while it was more highly expressed for MD *versus* MW at 10 DAP (Supplemental Data Set 2; row 100).

#### Structural genes in the phenylpropanoid biosynthesis pathway

In all, 109 genes were identified as putatively involved in the pathway of lignin biosynthesis based on gene annotation and orthologous relationships with genes from *G. max* and *A. thaliana* (Supplemental Data Set 3). No putative structural genes were identified within *qPD5.1-Pv*; however, several genes for lignin biosynthesis were located close to the major locus (Supplemental Figure 7). According to the RNA-seq expression data here, 50 (46%) of the total 109 structural genes were significantly differentially expressed for Midas *versus* G12873, for at least one of the two developmental stages that were considered (p <0.01; with 41 of these at p <0.001) (Supplemental Data Set 3). This suggests that the developmental phase between 5 DAP and 10 DAP is of particular importance for pod lignin biosynthesis.

#### *Expression patterns (qRT-PCR) of target candidates within the major locus* qPD5.1-Pv

The expression pattern for *PvMYB26* (Phvul.005G157600) was investigated in the pods of the totally indehiscent variety Midas, as well as for the three near isogenic ILs 038B/2A2, 244A/1A and 232B across eight pod developmental stages, using qRT-PCR. Up to the 4 DAP stage, no differential expression was seen between the mean expression of the three highly shattering lines and the totally indehiscent Midas (Figure 5).

**Figure 5.**
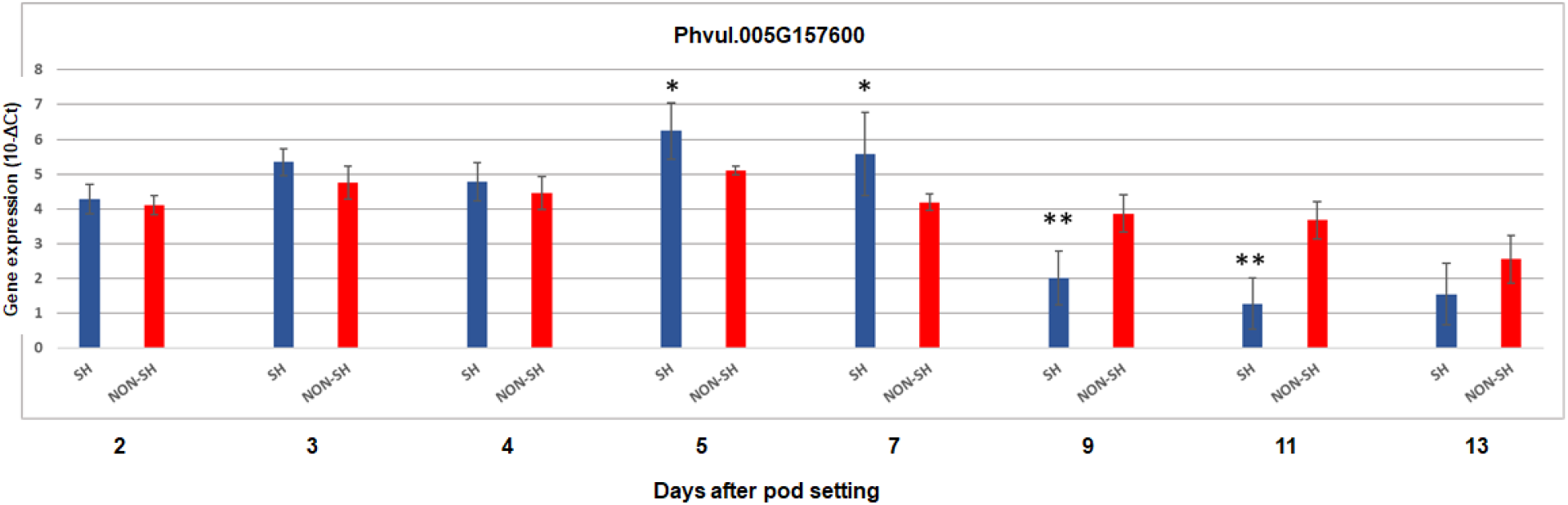
Gene expression by qRT-PCR for Phvul.005G157600 for the pods of the combined three highly dehiscent lines (SH; blue) and for the indehiscent pods of variety Midas (NON-SH; red) across the eight developmental stages from 2 DAP to 13 DAP. The mean pod expression for the three highly dehiscent introgression lines (038B/2A2, 244A/1A, 232B) is shown. *, p <0.05; **, p <0.01; SH *versus* NON-SH. Data are means ±standard deviation of the biological replicates (n = 3 for each highly dehiscent line for a total of nine for SH; n=4 for NON-SH). T.test for detection of significant differences, homoscedastic, two tails.

*PvMYB26* (Phvul.005G157600) was up-regulated at 5 DAP and 7 DAP in the dehiscent pods (fold-change, 2.20, 2.62, respectively; Supplemental Table 6), although at 7 DAP, only the expression of IL 232B was significantly different from Midas (Supplemental Figure 8). At 9 DAP, and with greater differences seen also at 11 DAP, *PvMYB26* was more highly expressed in the indehiscent pods of the variety Midas, as compared to the dehiscent lines, both as their combined mean expression (Figure 5) and as their individual expression (Supplemental Figure 8; and Supplemental Table 6). Reassuringly, the expression patterns for *PvMYB26* (Phvul.005G157600) were in agreement between the RNA-seq data (Midas *vs* G12873, Supplemental Data Set 1; row 50) and the qRT-PCR data. Among the target candidates for the major locus, efficient amplification was obtained for: Phvul.005G156900 (linoleate 9S-lipoxygenase); Phvul.005G161600 (translation initiation factor 2 subunit 3); Phvul.005G161800 (rRNA [uracil(747)-C(5)]-methyltransferase); Phvul.005G161900 (BHLH87 transcription factor similar to *AtIND*); Phvul.005G163901 (LEUCINE-RICH REPEAT PROTEIN KINASE-RELATED); Phvul.005G164800 (ZINC FINGER FYVE DOMAIN CONTAINING PROTEIN); Phvul.005G165600 (auxin-responsive protein IAA18-related); Phvul.005G165900 (LYSM domain receptor-like kinase); and Phvul.005G166300 (Myb-like DNA-binding domain). Phvul.005G161900 showed overall lower expression across the pod developmental stages and plant genotypes (for both qRT-PCR and RNA-seq) when compared to the other target candidates. However, slightly, but significantly, increased expression was seen for the dehiscent pods at 5 DAP (Supplemental Table 6).

As mentioned above, Phvul.005G156900 is a promising target candidate due to its genomic position and expression pattern (i.e., RNA-seq data). However, differential expression was observed only at 7 DAP for each of the dehiscent lines individually, but with variable expression patterns across the three dehiscent lines (Supplemental Table 6). Phvul.005G161800 showed higher expression in the dehiscent pods across all of the pod stages, with the greatest fold-change (3.273) seen for 11 DAP (Supplemental Table 6). These qRT-PCR data suggest that Phvul.005G161800 has a shattering-related function. The LRR-protein kinase related gene Phvul.005G163901 was highly expressed in the dehiscent pods with the most consistent differences seen at 4 DAP and 13 DAP (Supplemental Table 6). However, its expression pattern differed to that for the RNA-seq data (Midas *vs* G12873, Supplemental Data Set 1; row 110). This can potentially be explained by its expression being modulated after the expression of other genes involved in pod shattering, and its function is indeed worth further investigation.

When the shattering lines were considered as a combined group, Phvul.005G165900 showed lower expression in the highly shattering pods at 9 DAP, 11 DAP and 13 DAP (Supplemental Table 6). Moreover, the Phvul.005G165900 expression pattern was in agreement with the RNA-seq expression data (Midas *vs* G12873 at 10 DAP; see Supplemental Data Set 1; row 130).

Overall, the best target candidate genes for *qPD5.1-Pv* are summarised in Supplemental Table 7.

## DISCUSSION

Our results confirm that pod indehiscence in snap beans is controlled by a Mendelian locus with recessive inheritance. Here, we narrowed the major QTL *qPD5.1-Pv* down to a 22.5-kb genomic region that is located ~11 kb upstream of *PvMYB26*. Among the candidate genes for loss of pod shattering, *PvMYB26* is the best candidate because of its specific differential expression pattern between dehiscent and indehiscent pods, which is in agreement with the histological modifications associated to pod shattering across the same pod developmental phases. Moreover, the histological modifications are coherent with the function of *AtMYB26* in *A. thaliana*. Here, we also provide a list of candidate genes potentially involved in pod-shattering-related functions, through orthologue identification, selection signatures, and differential gene expression between wild and domesticated pods (i.e., RNA-seq) and/or between near isogenic lines (i.e., qRT-PCR).

We also demonstrate that pod indehiscence is associated with a lack of a functional abscission layer in the ventral sheath, due to ectopic lignification of a few layers of cells. Also, the key phenotypic events associated with pod shattering arise early in pod development, between 6 DAP and 10 DAP.

### Phenotypic architecture of pod shattering

Here, we propose that the failure of the formation of the abscission layer due to ectopic lignification is associated with pod indehiscence. This is similar to the ‘welding’ mechanisms previously defined for soybean by Dong *et al.* (2014), and more recently reported by Takahashi et al. (2019a) in an EMS mutant of *Vigna stipulacea*. Moreover, the cell-wall thickening pattern that we observed in the cells surrounding the abscission zone of the pods is in agreement to previous studies on *A. thaliana*, where in the wild-type, lignification at the valve margin close to the abscission layer is required for silique shattering (Liljegren *et al.*, 2004). Interestingly, valve margin lignification is also associated with pod coiling in *Medicago truncatula* (Fourquin *et al.*, 2013). We have also confirmed that an internal lignified valve layer forms in highly dehiscent pods, compared to indehiscent pods, which occurs early, before 10 DAP (Murgia *et al.*, 2017). This phenotype is most likely associated to the modulation of pod twisting, which from the phenotypic segregation analysis here, appears to be regulated in shattering pods by the action of at least two independent loci. Lignin deposition in the sclerenchyma of pod valves that is mediated by *GmPDH1* is also required for dehiscence modulation in soybean (Funatsuki *et al.*, 2014). This parallelism further reinforces the occurrence of convergent phenotypic evolution at the histological level between common bean and soybean for loss and reduction of pod shattering. Similarly, in some Brassicaceae, such as *Cardamine hirsuta*, asymmetric lignin deposition in *endocarp b* of the silique valves also ensures explosive seed dispersal and silique coiling (Hofhuis *et al.*, 2016). Furthermore, we propose that the key histological modifications associated to pod shattering occur between 6 DAP and 10 DAP. This agrees with the observation that 46% of the putative structural genes of lignin biosynthesis are differentially expressed in the same phase comparing indehiscent and highly shattering pods.

### *PvMYB26*: the best candidate for the major locus *qPD5.1-Pv*

Among the candidate genes that we investigated, we propose *PvMYB26* as the best candidate at the major locus for pod indehiscence. This is based on its genomic location, and on the parallel analysis of its expression patterns between dehiscent and indehiscent pods and of the histological modifications associated to pod shattering in the early phase of pod development. A role for *PvMYB26* in the loss of pod shattering is strongly supported also by the function of its orthologue in *A. thaliana*. Indeed, *AtMYB26* is required to establish which cells undergo cell-wall thickening to promote anther dehiscence (Yang et al., 2007 and 2017), and it acts upstream of the *NST1* and *NST2* genes, which have key roles in silique shattering (Mitsuda and Ohme-Takagi 2008). Interestingly, Takahashi *et al.* (2019b) suggested that pod shattering and pod tenderness are associated to MYB26 orthologues in azuki bean (*Vigna angularis*) and cowpea (*Vigna unguiculata*). The parallel identification of the MYB26 orthologue as the best candidate gene in *P. vulgaris* and other legumes (Takahashi *et al.*, 2019b), in addition to previous data from Rau *et al.* (2019) and Lo *et al.* (2018) in common bean and cowpea, respectively, further reinforce the hypothesis of the occurrence of molecular convergent evolution for domestication of pod shattering.

In addition to *PvMYB26*, we identified other genes that are worth highlighting. A cluster of four lipoxygenase genes were identified here, and their orthologues in *A. thaliana* (AT1G55020, AT3G22400) are putatively involved in defence responses, jasmonic acid biosynthesis, and responses to abscisic and jasmonic acid (The Arabidopsis Information Resource [TAIR] database). We also highlight Phvul.005G163901 and Phvul.005G163800 within a cluster of LRR genes, and Phvul.005G161800 (rRNA [uracil(747)-C(5)]-methyltransferase). Interestingly, a potential role for LRR-RLK genes in shattering-related functions, such as secondary cell-wall biosynthesis and abscission processes, can be postulated according to Jinn *et al.* (2000), Bryan *et al.* (2011), Van Der Does *et al.* (2017) and Xu *et al.* (2008).

Overall, no putative structural genes for lignin fell within *qPD5.1-Pv*. We suggest that selection might preferentially act on regulation factors instead of genes with a central role in the lignin biosynthetic pathway, perturbations of which can result in side-effects on genotype fitness and/or can be disabling for normal development of the plant. However, there was a cluster of putative structural genes for lignin biosynthesis close to *qPD5.1-Pv*, which suggests that they are directly involved in the same pathway of the genes responsible for the major QTL.

Based on the evidence we present here, *PvMYB26* is the best candidate for the major locus. Nevertheless, the presence of further candidates that are also organised within a cluster of genes suggests that the main QTL operates in an ‘operon’-like manner. Indeed, the clustering of duplicated or non-orthologous genes might provide advantages in terms of coordination of expression between physically close genes that are involved in the same pathway, as for secondary metabolite biosynthesis (Osbourn 2010 and Boycheva *et al.*, 2014).

### Convergent evolution and conservation of the molecular pathway for modulation of pod shattering

In the present study, we identified orthologous genes that are putatively involved in pod-shattering-related functions, and we analysed their expression profiles and selection signatures to provide potential candidates at known QTLs and/or at the genome-wide level. Among these, we highlight Phvul.002G271000 (*PvIND*), as orthologous to *AtIND* (Liljegren *et al.*, 2004), which has a pivotal role in silique shattering in *A. thaliana*. Moreover, our expression data and selection signatures reinforce the orthologue of *PDH1* (Funatsuki *et al.*, 2014) (*PvPdh1*; Phvul.003G252100) as a strong candidate for modulation of pod shattering also in common bean (Parker *et al.*, 2020). This might further suggest the occurrence of selection at orthologous loci for loss or reduction of pod shattering between closely related legume species. In addition, Phvul.009G203400 is a promising target candidate that shows parallel selection across the Andean and Mesoamerican gene pools, according to both Schmutz *et al.* (2014) and Bellucci *et al.* (2014). Moreover, Phvul.009G203400 is the orthologous to *AtFUL* (Gu *et al.*, 1998) which is involved in valve differentiation in *A. thaliana*. Here, we also identified Phvul.010G118700, as orthologous to *NST1* (Mitsuda and Ohme-Takagi 2008) and *GmSHAT1-5* (Dong *et al.*, 2014), which have crucial roles in silique shattering and in pod shattering resistance in *A. thaliana* and soybean, respectively. In addition to the major candidate *PvMYB26*, we also identified several MYB-like protein-coding genes close to known QTLs or at the genome-wide level (see Supplemental Data Set 2), and among these, a paralogue to *PvMYB26* on chromosome Pv10. The function of MYB transcription factors in the regulation of both secondary cell-wall biosynthesis and the phenylpropanoid pathway has been widely reported (Zhong *et al.*, 2008, Zhang *et al.*, 2018). Overall, the expression patterns between the wild and domesticated pods, and the presence of selection signatures at orthologous genes at the genome-wide level (see Supplemental Data Set 2), suggest that several of these have preserved shattering-related functions, and that there has been conservation across distant taxa of the pathway associated to seed dispersal mechanisms. This was previously demonstrated in rice (Konishi *et al.*, 2006, Yoon *et al*., 2014), soybean (Dong *et al.*, 2014) and tomato (Vrebalov *et al.*, 2009).

Identification of the molecular and phenotypic bases of a key trait of domestication in common bean such as pod shattering represents a fundamental step in the dissecting out of the evolutionary history of the species. On this basis, we proposed here *PvMYB26* at the major locus that controls pod indehiscence, and we have illustrated the key histological changes at the pod level that are associated with pod shattering. We also used a strategy that combined expression analysis, signature selection and orthologous identification, which allowed us to identify a number of potential target candidate genes that need to be validated in future studies. We believe that the increasing number of studies on the architecture of pod shattering will allow reconstruction of the evolutionary trajectories that have led to convergent modifications across legumes with ever greater resolution. With this perspective, and based on the data we have presented here, we propose that loss and reduction of pod shattering might arise after selection at orthologous loci, and that this has resulted in convergent evolution also at the histological level.

## Materials and Methods

### Development of the IL population

Here, we developed an IL population (1,197 BC4/F4) for the mapping of pod shattering traits (see Supplemental Figure 9). The IL population was developed starting from a cross between the domesticated Andean variety Midas, as ‘stringless’ and totally indehiscent, and the highly shattering wild Mesoamerican genotype G12873, to provide an initial set of RILs (Koinange *et al*., 1996). One RIL (i.e., MG38) showed high shattering, wild traits of the seeds and pods, a determinate growth habit, and the absence of photoperiod sensitivity, so it was selected as a donor parent for pod-shattering traits for backcrossing with the recurrent Midas (BC1). Overall, three backcrosses were performed using Midas as the recurrent parent and maintaining the phenotypic selection for high shattering for each backcrossed generation, which provided 70 ILs from BC3/F4:F5 families, and 217 ILs from BC3/F6:F7 families (Murgia *et al.*, 2017, Rau *et al.*, 2019). In the present study, six highly shattering ILs were selected as the donor parents for high pod shattering, and were further backcrossed (BC4) with Midas, providing six subpopulations (BC4/F1 families), for the lines: 232B (from a BC3/F4:F5 family); and 244A/1A, 038B/2A2, 038B/2C1, 038A/2D1 and 038B/2B1 (from BC3/F6:F7 families). Seeds of BC4/F1 individuals and of the seven parental lines were pre-germinated in Petri dishes using deionised water. The plants were individually grown in the greenhouse of the *Dipartimento di Scienze Agrarie, Alimentari ed Ambientali* at the Polytechnic University of Marche in Ancona, Italy, between January and May 2016. BC4/F2 seeds were collected from 100 BC4/F1 lines, and 1,353 BC4/F2 harvested seeds were planted in open field at Villa D’Agri, Marsicovetere (Potenza, Italy), in the summer of 2016. Some of these (636 BC4/F2 seeds) were pre-germinated using vermiculite and deionised water, and the seedlings were transplanted on the first planting (7 June 2016), while the other 717 BC4/F2 seeds were directly sown as a second planting (26 July 2016). The pods were collected from 942 BC4/F2:F3 ILs in October 2016. The BC4/F3 plants were obtained by single seed descent and grown in the greenhouse between February and May 2017. With the aim to reach an initial population size of 1,000 BC4/F3 individuals, two BC4/F3 seeds were sown from a few dehiscent BC4/F2 lines. The pods were collected from 724 BC4/F3 individuals. Then 2,230 BC4/F4 seeds and 109 seeds from the seven parental lines of the new population were sown in an open field at Villa D’Agri in the summer of 2017. The seeds were directly sown on 22^nd^ June, and additional sowing was performed to any recover missing plants. One BC4/F4 seed from each BC4/F3 indehiscent line, and at least four BC4/F4 seeds from each BC4/F3 dehiscent line were sown, with the aim to promote segregation and recombination at the major locus *qPD5.1-Pv* for pod indehiscence on Pv05 (Rau *et al.*, 2019), at which a recessive domesticated allele determines the totally indehiscent phenotype only in the homozygous condition. The pods were collected from 1,197 BC4/F4:F5 ILs. The BC4/F2 experimental field scheme provided 12 rows, with sowing distance of 0.6 m and 1.5 m within and between the rows, respectively. The BC4/F4 field scheme consisted of 2,339 holes across 9 rows, with sowing distance of 0.25 m and 1.2 m within and between the rows, respectively. In the field trials, the ILs were completely randomised within the six BC4/F1 families. Weed control was provided using a mulching plastic sheet, and pest control treatments were with Ridomil Gold (fungicide) and Klartan 20 Ew (against aphids). The plants were watered daily using an automatic irrigation system, and fertilisation with nitrogen, phosphorous and potassium was applied before tillage.

### Phenotyping of the IL population

Phenotyping for pod shattering was performed in the field trials both qualitatively (i.e., occurrence of pod shattering, with each plant classified as dehiscent if it showed at least one shattered pod), and quantitatively, by assigning a score to each dehiscent line based on the pod twisting: 0 (no twisted pods per plant); 1 (1% < twisted pods < 10%); 2 (≥10% < twisted pods < 24%); and 3 (≥24% of twisted pods). Shattering was evaluated in the BC4/F2 ILs across four dates (19 September, 29 September, 13 October, 23 October) until the uniform ripening of the entire plants, and in the BC4/F4 lines across two main dates (18 October, 22 October), plus two additional dates (26 October, 12 November) for plants which were not fully ripened at the earlier dates. Pod shattering was also evaluated post-harvest by examination of the completely dry pods. For the BC4/F1 individuals, each genotype was classified as easy to thresh (i.e., pods opened very easy along sutures), similar to the highly shattering parents, or as totally indehiscent, similar to the domesticated parent Midas. For the other experiments, phenotyping was performed by testing the resistance to opening when the ripened pods were subjected to increasing manual pressure directly on the sutures, according to the scoring system in Supplemental Table 8. Moreover, a comprehensive phenotypic trait for pod shattering was assigned manually to each BC4/F4 line (i.e., ‘SH y/n’; presence or absence of pod dehiscence), which combined field and post-harvest phenotyping. A few BC/F4 ILs were classified as intermediate when it was not possible to assign an accurate phenotype to the line.

### Genotyping and genome-wide association study for pod shattering

Young leaves were collected from 1,197 BC4/F4 ILs and 55 replicates from the seven parental lines that were grown during the last IL field experiment. The leaves were dried within 12 h of collection using silica gel. Genomic DNA (gDNA) was extracted from the leaves using the Exgene Plant SV kit (Geneall Biotechnology) and stored at −20 °C. The gDNA integrity was determined on 1% agarose gels, and the DNA quality was measured using a photometer (NanoPhotometer NP80; Implen) and quantified with the dsDNA assay kits (Qubit HS; Life Technologies). The gDNA concentrations were adjusted to 25 ng/μL, and the genotyping was performed using GBS (Elshire *et al.*, 2011) by Personal Genomics (Verona, Italy). The protocol for the GBS library preparation is provided in the Supplemental Methods (GBS library preparation), according to the procedure reported in Rau et *al*. 2019. The GBS libraries were sequenced (HiSeqX platform; Illumina with 2x 150 bp reads mode at Macrogen Inc. [South Korea]), which generated 1.5 million fragments per sample on average. The sequencing reads were demultiplexed based on their barcodes. Adapters and low-quality bases in the FASTQ files were removed using the Cutadapt software, version 1.8.3 (Martin 2011). The filtered reads were aligned to the reference genome of *P. vulgaris* 442 version 2.0 using the BWA-mem software, version 0.7.17-r1188 (Li and Durbin 2009). The resulting BAM files were realigned using the GATK RealignerTargetCreator and IndelRealigner software, version 3.8.1, to remove errors. Variant calling was performed for all of the samples together, using the GATK UnifiedGenotyper software, version 3.8.1 (McKenna *et al*., 2010), and the variants were filtered based on GATK best practice. The raw SNP dataset (2,419,927 SNPs) was checked for quality and loci with missing data >95% and with MAF < 0.05 were excluded from further analysis. Additionally, filtering was performed to remove SNPs that were either missing in one parental set (i.e. MIDAS or MG38), monomorphic between parents or located in SCAFFOLDS (as SCAFFOLDS resulted not associated any of the investigated traits). The dataset was then imputed for missing data by using beagle.5 (Browning *et al*., 2018). A further filtering was performed after imputation to remove few more sites that were monomorphic between the parents. The final data set included 1253 individuals (i.e. 1196 BC/F4 ILs, 55 parental lines of the BC4 population, and the references Midas and MG38) and 19,420 SNPs. GWAS was performed by using the Mixed Linear Model (MLM) as implemented in the rMVP package (https://github.com/xiaolei-lab/rMVP). Overall, seven descriptors of pod shattering were considered for GWAS analysis from the three main phenotypic dataset (i.e., Sh y/n [integration of field and post-harvest data], field, and post-harvest): Sh y/n (dehiscent *vs* indehiscent lines), Sh y/n after filtering (18 lines that showed signs of diseases and/or a low pod production were removed), Sh y/n including lines with an intermediate phenotype between the dehiscent and the indehiscent, Field (presence *vs* absence of pod shattering), Field (percentage of twisting pods per plant), Post-harvest (putative dehiscent *vs* putative indehiscent), Post-harvest (quantititative; mapping of all the phenotypic classes 0, 1, 1.5, 2, 3).

### RNA sequencing and differential gene expression analysis

The wild dehiscent Mesoamerican genotype G12873 and the fully indehiscent Andean variety Midas were grown for the collection of their pods under controlled conditions in a growth chamber at the Institute of Biosciences and Geosciences (IBG-2, Forschungszentrum Jülich), in 2014. Two plants were planted for both the G12873 and Midas genotypes. In the same experiment, a total of 57 plants were grown from 43 different genotypes, as for 14 of these, two replicates were available. Considering the overall number of plants, nine were AD, 18 were AW, 12 were MD, and 18 were MW. Moreover, three of the nine AD were snap bean types (AD_Snap; totally indehiscent), while the other six were dry beans, according to the phenotypic data and the available information. The list of the accessions is provided in Supplemental Data Set 4. The experimental conditions were 24/20 °C day/night temperature, 70% of relative air humidity, photon lux density of 400-500 μmol m-2 s-1, and short-day photoperiod conditions (10/14 h light/dark). Fertilization was provided for N-K-P and trace elements. The pods were collected for each genotype at 5 DAP and 10 DAP. These were snap-frozen in liquid nitrogen before storage at –80 °C. After RNA extraction, the cDNA libraries were prepared according to the Illumina TruSeq RNA LT protocol, and the RNA sequencing was performed with the HiSeq paired-end V4/4000 125/150 cycles sequencing technology. Sequencing was performed by the Genomics and Microarray Core Laboratory at the University of Colorado in Denver (USA), and the raw-data quality check and alignment were performed by Sequentia Biotech (Barcelona, Spain). The read quality checking was performed on the raw sequencing data using the FastQC tool, and low-quality portions of the reads were removed using BBduk. The minimum length of the reads after trimming was set to 35 bp, and the minimum base quality score to 25. High quality reads were aligned against the *P. vulgaris* reference genome (G19833 genome v2.1; http://phytozome.jgi.doe.gov/) using the STAR aligner software, version 2.5.0c. The reads that could not be aligned against the first reference genome were mapped against the second reference genome (*P. vulgaris* L., BAT93; [Vlasova *et al.*, 2016]). FeatureCounts, version 1.4.6-p5, was used to calculate the gene expression values as raw read counts. Here, the raw reads data were used to perform the differential gene expression analysis across the two developmental stages, using the DESeq2 package (Love *et al.*, 2014) in R (R Core Team 2019). The differential gene expression was calculated for each gene (as log_2_ fold-changes), and the p values were adjusted according to the Benjamini-Hochberg procedure (Benjamini and Hochberg 1995). Differential gene expression was performed at 5 DAP and 10 DAP for the following comparisons: Midas *versus* G12873, AD *versus* AW; AD_Snap *versus* AD; AD_Snap *versus* AW; and MD *versus* MW.

### *qRT-PCR of candidate genes for the* qPD5.1-Pv *locus*

The indehiscent variety Midas and three parental lines of the IL mapping population with the highest level of pod shattering (ILs 232B, 244A/1A, 038B/2A2), and that were near isogenic to Midas after three backcrosses, were grown in a greenhouse at the Max Planck Institute of Molecular Plant Physiology (Golm-Potsdam, Germany), in April to July 2018. The plants were individually grown in pots (diameter, 20 cm; volume, 3 L), and fertilisation was performed with Hakaphos rot (0.015%) during irrigation (666 g/10 L). The plants were watered four times per day, and pest control was performed using Neem Azal (6 mL/3 L). At least nine biological replicates were grown for each genotype. At least three pods from each dehiscent genotype and four pods for Midas were collected from different replicates, at 2, 3, 4, 5, 7, 9, 11 and 13 DAP. Entire green pods were collected from 2 DAP to 5 DAP, while from 7 DAP, the ventral and dorsal sutures were separated manually from the valves and collected separately to evaluate gene expression in the region surrounding the ventral suture. The pods were frozen in liquid nitrogen before storage at −80 °C. The pod tissues were ground with a mixer mill (MM400; Retsch), and the RNA was extracted using the RNA miniprep kit (Direct-zol; Zymo Research GmbH). The RNA was stained using GelRed, and its integrity was visualised using 1% agarose gels. The RNA concentrations and quality were measured using a spectrophotometer (NanoDrop OneC; Thermo Scientific). After adjusting the RNA concentrations, the cDNA was synthesized for each sample (Maxima First Strand cDNA Synthesis Kit with dsDNase; Thermo Scientific). Each cDNA was diluted 1:10, by adding HPLC quality water, and stored at −80 °C. The primers for the candidate genes (i.e., qRT-PCR) were designed based on the gene coding sequences using the Primer3 (v0.4.0) tool (Supplemental Table 9). The target candidate genes were selected based on gene annotation, gene expression from the RNA-seq data, the presence of selection signatures according to Schmutz *et al.* (2014) and Bellucci *et al.* (2014), the functions of orthologous genes, and the location in the genomic regions with high association to pod shattering. Two housekeeping genes were included, based on the literature (i.e., Phvul.007G270100 (Borges *et al.*, 2012); Phvul.010G122200 (Montero-Tavera *et al.*, 2017). The amplification efficiencies were determined for each pair of primers. Here, four dilutions (i.e., 1:10, 1:20, 1:30, 1:40) of the same cDNA were amplified (i.e., qRT-PCR), and the slope (R^2^) of the calibration curve was used to infer the primer efficiency, according to Equation (1):

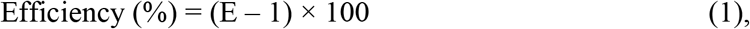

where E was obtained from R^2^ according to the Equation (2):

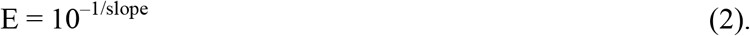

The differential gene expression was calculated as fold-changes between each dehiscent line (i.e., 232B, 244A/1A, 038B/2A2) and the indehiscent line Midas, and for all of the donor parents grouped together *versus* Midas, according to Schmittgen and Livak (2008). T-tests were performed for each comparison separately, as comparisons of the ΔCt values. ΔCt was obtained as the difference between the Ct (cycle threshold) of the candidate gene and the Ct of the housekeeping gene for normalisation of gene expression, according to Schmittgen and Livak (2008).

### Identification of orthologous genes with putative functions in pod shattering

The common bean genome (v2.1) (http://phytozome.jgi.doe.gov/) contains 27,433 loci and 36,995 protein-coding transcripts. However, for several of these, the annotation is missing or not always accurate. We used the Orthofinder algorithm (Emms and Kelly 2015) to identify clusters of orthologous genes among the proteome of *P. vulgaris*, nine related legume species and *A. thaliana*. The proteome sequences considered here were: *A. thaliana* (TAIR10); *P. vulgaris* (v2.1); *G. max* (Wm82.a2.v1); *M. truncatula* (285_Mt4.0v1); *V. unguiculata* (v1.1); *Cicer arietinum* (cicar.ICC4958.gnm2.ann1); *Lotus japonicus* (v3.0); *Lupinus angustifolius* (1.0); *Vigna angularis* (vigan.Gyeongwon.gnm3.ann1.3Nz5); *Vigna radiata* (vigra.VC1973A.gnm6.ann1); and *Glycyrrhiza uralensis* (Gur.draft-genome.20151208). These were downloaded from: Phytozome (http://phytozome.jgi.doe.gov/); the ILS database (https://legumeinfo.org/); the *Lotus japonicus* genome assembly (http://www.kazusa.or.jp/lotus/); and the *Glycyrrhiza uralensis* genome database (http://ngs-data-archive.psc.riken.jp/Gur-genome/index.pl). The protein sequences from the primary transcripts were used for the analysis, except for *L. japonicus*, for which only the full proteome was available. Orthofinder (v2.1.2) identified 20,692 orthogroups (i.e., clusters of orthologous genes) across these 11 species. The list of structural genes involved in the synthesis of phenylpropanoid was obtained from the Plant Metabolic Network database (https://www.plantcyc.org/) for common bean, soybean and *A. thaliana*, as pod shattering in common bean is positively associated with lignin content in the pods (Murgia *et al.*, 2017). Common bean genes without any clear annotation were considered as putative structural genes of phenylpropanoid synthesis if they clustered in the same orthogroup of *A. thaliana* and soybean lignin biosynthesis-related genes. *A. thaliana* and soybean lignin-related genes that were not assigned to any orthogroup were blasted with BLASTp against the common bean proteome to identify the best putative orthologues. Common bean gene orthologues to those with a well-known role or a putative function in seed dispersal mechanisms in *A. thaliana* and in other crops, according to the available literature, were also identified with the same approach.

### Identification of selection signatures

Genes that underwent selection during domestication of common bean in Mesoamerica and in the Andes (Schmutz *et al.*, 2014) were identified. Moreover, 27,243 contigs that were previously detected by Bellucci *et al.* (2014), which included 2,364 putatively under selection in the Mesoamerican pool, were mapped against the last common bean genome version. Briefly, the contigs were aligned against the *P. vulgaris* protein sequences of all of the gene coding sequences (annotation on Phytozome, version 2.1) using NCBI blastx (blast-2.2.26), and then the best hit for each contig was selected and the reference gene of each contig was established with a threshold of <1 E-10. A gene was considered as putatively under selection if at least one of the five contigs with the best e values was putatively under selection in Bellucci *et al.* (2014).

### Pod histological analysis on parental lines of the IL population

Pods of the highly shattering genotypes 232B, 244A/1A and 038B/2A2 (ILs) and the totally indehiscent variety Midas were collected for histological investigation. These were from the same greenhouse experiment that was performed for the qRT-PCR expression analysis. In addition, replicates of genotype MG38 (RIL) were grown in the same experiment. Entire pods were collected across five developmental stages (6, 10, 14, 18, 30 DAP). Then 2-cm to 3-cm free-hand cross sections from the pods were fixed in 5% agarose, and 70 μm, 50 μm and 30 μm cross-sections were obtained using a microtome (VT 1000 S; Leica). A solution of phloroglucinol (7 mg), methanol (7 mL) and 37% chloridric acid (7 mL) was applied to the pod sections for specific lignin staining. The pod sections were visualised under an optical microscope (BX51TF; Olympus).

## Supplemental Data

**Supplemental Figure 1**. Analysis of lignification patterns in the dorsal sheaths of 6-day-old pods of the totally indehiscent variety Midas (A, B) and the highly pod shattering IL 244A/1A (C, D).

**Supplemental Figure 2**. Analysis of lignification patterns in pod valves of 10-day-old pods of the totally indehiscent variety Midas (A), and of two highly pod shattering RIL MG38 (B) and IL 244A/1A(C).

**Supplemental Figure 3**. Analysis of lignification patterns in the ventral sheaths of 14-day-old pods of the totally indehiscent variety Midas (A, B) and the highly pod shattering RIL MG38 (C, D) and IL 244A/1A (E, F).

**Supplemental Figure 4**. Densities of the 19,420 SNP markers identified within a 1-Mb window size using genotyping by sequencing.

**Supplemental Figure 5**. Genome-wide association study for occurrence of pod shattering on the IL population.

**Supplemental Figure 6**. Expanded major QTL for pod shattering on chromosome Pv05.

**Supplemental Figure 7**. Physical positions of the putative structural genes for lignin biosynthesis on the common bean chromosomes.

**Supplemental Figure 8**. Gene expression by qRT-PCR for Phvul.005G157600 for the pods of the three highly dehiscent ILs (as indicated, blue) and for the indehiscent pods of variety Midas (MIDAS, red) across the eight developmental stages from 2 DAP to 13 DAP.

**Supplemental Figure 9**. Schematic representation of the development of the BC4/F4 introgression line population.

**Supplemental Figure 10**. Structure of the GBS library.

**Supplemental Table 1**. Segregation of pod shattering on a subset of the BC4/F2 lines.

**Supplemental Table 2**. Observed segregation for the trait of ‘pod shattering occurrence’ in the BC4/F2 population, and for each subpopulation.

**Supplemental Table 3**. Results of the post-harvest phenotyping for pod shattering for 549 BC4/F3 Ils.

**Supplemental Table 4**. Results of field and post-harvest phenotyping for pod shattering for 1,197 BC4/F4 ILs.

**Supplemental Table 5**. Summary of the genome-wide association study for pod shattering in the BC/F4 ILs population.

**Supplemental Table 6**. Differential gene expression by qRT-PCR of the target candidate genes at the major locus *qPD5.1-Pv* for pod indehiscence.

**Supplemental Table 7**. Summary of the best candidate genes at the major locus *qPD5.1-Pv*, according to the expression data (i.e., RNA-seq, qRT-PCR), the presence of a selection signature (Schmutz *et al.* (2014) and Bellucci *et al.* [20]), and gene annotation of the common bean gene and its orthologues in *A. thaliana* and other crops.

**Supplemental Table 8**. Post-harvest phenotyping for the scoring of pod shattering of the IL population.

**Supplemental Table 9**. Primers sequences for qRT-PCR and gene expression analysis of the target candidate genes at the major locus *qPD5.1-Pv* for pod indehiscence.

**Supplemental Table 10**. Sequences of the single-stranded oligos for the adapters used for GBS library preparation.

**Supplemental Table 11**. Sequences of the primers used for the amplification, indexing and quantification of the GBS library.

**Supplemental Data Set 1**. Genes identified within the major locus *qPD5.1-Pv* for loss of pod shattering.

**Supplemental Data Set 2**. Genes in common bean that are orthologous to genes in other species with known functions that are putatively involved in seed shattering or have potentially related functions (e.g., cell-wall modification, differentiation).

**Supplemental Data Set 3**. Genes in common bean that are putatively involved in the phenylpropanoid biosynthesis pathway.

**Supplemental Data Set 4**. List of accessions that were grown for pod collection, RNA-seq and differential gene expression analyses.

## SUPPLEMENTAL METHODS

*GBS library preparation*

## SUPPLEMENTAL REFERENCES

## Acknowledgments

The study was part of the PhD research work of the first author (V.D) and he is grateful to the Doctoral School of the Polytechnic University of Marche (UNIVPM) and to the FORSCHUNGSZENTRUM JÜLICH (IBG-2) that co-funded the PhD scholarship.

This work was supported by the BRESOV project, funded from the European Union’s Horizon 2020 research and innovation program under Grant Agreement No. 774244, the Italian Government (MIUR; Grant number RBFR13IDFM_001, FIRB Project 2013), and the Polytechnic University of Marche.

Alisdair R. Fernie and Saleh Alseekh acknowledge the European Union project PlantaSyst (SGA-CSA No 664621 and No 739582 under FPA No. 664620).

We are grateful to Prof. Paul Gepts for providing the MG38 genotype, which was used as a donor parental line for the development of the population in the earlier stage of the research, and to the Sequentia Biotech for the bioinformatic analyses on RNA-seq data.

Valerio Di Vittori is grateful to Dr Leonardo Perez de Souza, José G. Vallarino and Federico Scossa for their help and critical input in the differential gene expression analysis and identification of orthologous genes.

## Author Contributions

**R.P** conceived and supervised the study; **V.D** and **R.P** designed the study and wrote the article; **V.D**, **E.Bi**, **E.Be, L.N**, **D.R**, **M.Rod**, **GA**, **J.F**, **A.C, G.L, S.M** and **R.P** contributed in the development of the IL population; **V.D**, **T.G, M.L.M, F.F and B.U** were involved in the phenotyping; **V.D**, **M.Ros, C.D**, **M.D and C.X** were involved in genotyping and bioinformatic analyses; **M.Rod** performed GWAS; **E.B** provided the RNA-seq data; **V.D**. performed the qRT-PCR experiment, the differential expression analysis, the histological analysis and orthologue identification under the supervision of **S.A** and **A.R.F** and with critical input from **A.S** and **A.F**. All of the authors have read and approved the final version of the manuscript, with further critical input.

